# A rapid method for DNA extraction of cotton mature fiber suitable for PCR fingerprinting

**DOI:** 10.1101/529305

**Authors:** Shafik D. Ibrahim, Alsamman M. Alsamman, Kareem Khalifa

## Abstract

Egyptian cotton with its extra-long staple fiber is the world’s finest textile crop, but it has been in a state of rapid decline for the past 20 years. One of the main problems in Egyptian cotton production that, some manufacturers are using inferior quality cotton or other cheaper cotton to cut costs. This practice impacts not only the brand owner but also the cotton farmers and producers. The ability to differentiate between different mature or processed cottons fibers according to its varietal origin is very complicated or highly expensive. We introduce the first non commercial procedure to isolate intact genomic DNA from cotton fiber with a modified CTAB protocol. This protocol was successfully used to retrieve DNA from cotton fabric and the DNA was efficient to perform PCR procedure using DNA-barcoding and SCoT assays. Additionally, we retrieved PCR markers using CAPs assay can differentiate between Egyptian and American cotton from a fiber source.

## Introduction

With a $12 billion trade and 25 million tonnes production, cotton is the world’s most popular natural fiber and an important commodity in the world economy. More than two thirds of the world’s cotton is produced by developing countries, where it is an important cash crop at both farm and national levels, through their spinning and textile industries (Gillson 2004).

Egyptian cotton with its extra-long staple fiber is the world’s finest textile crop, but it has been in a state of rapid decline for the past 20 years. Regardless of the $0.5 billion support Egyptian government introduces yearly to cotton farmers; its production is predicted to drop by 40 percent for the upcoming 2015/2016 crop from 525,000 bales in the current marketing season and cotton planted area is expected to drop to 240000 feddans in 2016. One of the main problems in Egyptian cotton production that, some manufacturers are using inferior quality cotton or other cheaper cotton to cut costs. This practice impacts not only the brand owner but also the cotton farmers and producers. On the other hand, it costs Egypt millions or perhaps billions of Dollars annually through losing open global markets and customers.

The ability to differentiate between different mature or processed cottons fibers according to its varietal origin is very complicated or highly expensive. The first step to is to obtain a good quality DNA amounts efficient for different DNA-finger printing techniques. The second step is to use non-expensive and highly throughput methods to differentiate between different samples used in clothes industries without addition more charge on the clothes production in the final customer price.

Although there are some published studies that claim the ability to extract DNA from cotton fibers (US8940485 B2) it is unfortunately for non-public use and it must be paid for using which could avoid cotton researchers from breaking through this areas or cotton producers from using molecular genetics techniques as efficient methodology to avoiding commercial fraud. Being able to identify the cultivar of a particular species of cotton utilized in a textile item would not only be a way to authenticate an item as legitimate and being made with the type of cotton specified by the owners, but would also enable the detection of forged or counterfeit textile products.

The term “DNA barcode” for global species identification was first coined by Hebert et al., in 2003 and has gained worldwide attention in the scientific community. Recognition of animals, plants and fungi has been performed using this technique. Screening for single or multiple regions appropriate for DNA bar-coding studies in nuclear and plastid genomes in plants has been an important research focus.

CTAB DNA purification methods extract high quantities of pure DNA from a variety of different plant tissues. However, there is no protocols is suitable for extracting efficient-quality DNA from cotton fibers. In this report, we describe a simple and reliable CTAB method for isolation of amplifiable DNA with as little as 200 ng of cotton fiber materials. In addition we used DNA-barcoding and CAPs techniques to differentiate between Egyptian and imported cotton cultivars.

## Methodology

### DNA Isolation

Seven mature cotton fiber and one plant leaf samples have been used for DNA extraction, where four are Egyptian cotton cultivars, three American cultivars and one unknown lab cotton (cotton fibers use for lab and medical applications) (commercial).

All tools were sterilized using autoclave and all work was conducted under laminar flow conditions. About 0.2g of cotton fibers was disinfected in 100 distilled water with 10% bleach (commercial Clorox) for 10 minutes, then with 70% alcohol for 20 minutes and finally washed with deionized and distilled water. After drying, fibers were ground with mortar and pestle along with some glass cover slips (Cat No 7101, Sigma Aldrich) until all mix became fine powder (it could took 10 minutes for every sample). DNA Extraction was isolated from fiber powders and young leaf material following the CTAB procedure of Porebski et al. (1997).

### SCoT PCR assay

The PCR amplification reaction was carried out for one SCoT primer. The PCR was performed in 25 μl volume composed of 1x reaction buffer, 0.2 mM of dNTPs, 1.5 mM MgCl2, 0.2 μM of primer, 1U of *Taq* polymerase and 15ng of template DNA, in sterile distilled water. PCR amplification of the DNA was performed in a Perkin Elmer thermal cycler 9700. The temperature profile in the different cycles was as follows: an initial strand separation cycle at 94°C for 5 min followed by 40 cycles comprised of a denaturation step at 94°C for 40sec. an annealing step at 50°C for 50sec. and an extension step at 72°C for 1 min. The final cycle was a polymerization cycle for 7 min at 72°C. PCR products were resolved by electrophoresis in a 1.5% agarose gel containing ethidium bromide (0.5 mg/ml) in 1 x TBE buffer at 120 volts. A 100bp DNA ladder was used as molecular size standard. PCR products were visualized under UV light and documented using a ™XR+ Gel Documentation System (Bio-Rad).

### Rbcl DNA-Barcoding assay

DNA barcoding analysis was performed with the plastidial RbcL region. For PCR amplification and sequencing of RbcL, the primer combination was Forward primer: 5′-ATGTCACCACAAACAGAGACTAAAGC-3′and Reverse primer: 5′-TCGCATGTACCTGCAGTAGC-3′. The reaction mixture consisted of 1× buffer (Promega), 15mM MgCl2, 0.2 mM dNTPs, 20pcoml of each primers, 1 u of Taq DNA polymerase (GoTaq, Promega), 30 ng DNA and ultra-pure water to a final volume of 20 uL. The following PCR program was used: 94 °C for 2 min followed by 40 cycles at 94°C for 30 s, 55°C for 30 s, 72 °C for 60 s. with an additional cycle at 72 °C for 7 min. DNA Sanger sequencing was carried out by Macrogen inc., Korea. The NCBI online BLAST tool was used with its default parameters to align the generated sequences using BLAST algorithm (Altschul et al., 1990) against the NCBI database.

### Cleaved Amplified Polymorphic Sequences analysis

To recover polymorphism between monomorphic bands, restriction enzyme digestion was performed in order to develop CAPs (cleaved amplified polymorphic sequences) for restriction assays, 10 μl PCR products were incubated overnight at 37°C in a volume of 25 μl with 3 U of a restriction enzyme, and subsequently resolved in a 2.5% agarose gel. Restriction with 12 enzymes was assayed. Finally, HeaIII and BlgII (Fermentas, Life Sciences) were the enzymes used to generate polymorphisms.

## Results and Discussion

The quantity and quality of extracted genomic DNA from different amounts of mature cotton fibers is low but it was very efficient to perform all PCR procedures (SCoT and DNA-barcoding). On the other hand, this DNA isolation technique was very time-saving and it did not take any expensive procedures.

Successful PCR amplification of the chloroplast/chromoplast RbcL gene promoter gene and SCoT markers indicates that organelles’ genomic DNA was co-extracted with nuclear genomic DNA and both were of high quality and amplifiable (Figures 1 and 2).

**Figure 1:**
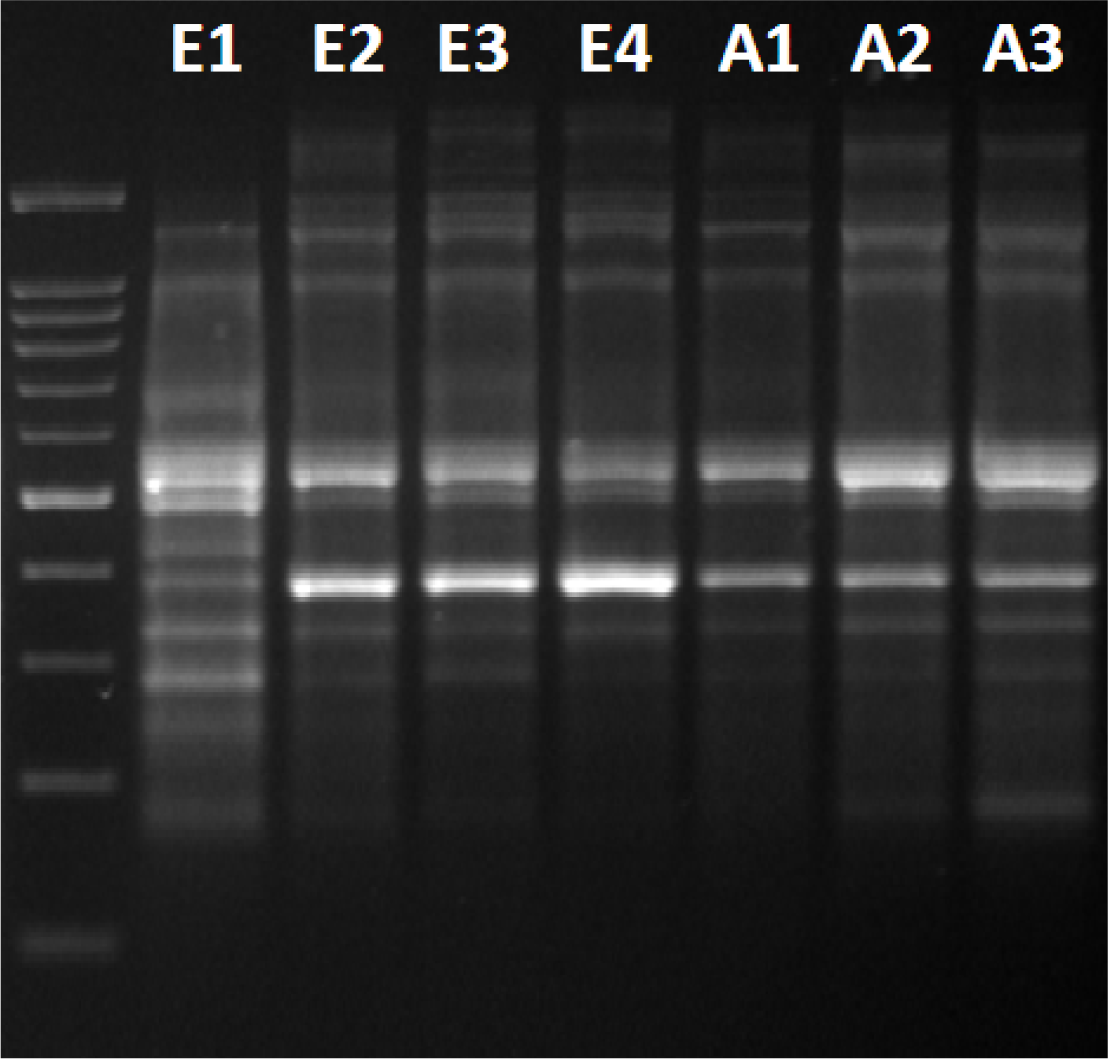
The PCR products of SCoT assay for the DNA isolated form Egyptian cotton plant leaf (E1), Egyptian cotton fiber (E2, E3, E4) and American cotton fiber (E5,E6, E7) varieties.

**Figure 2:**
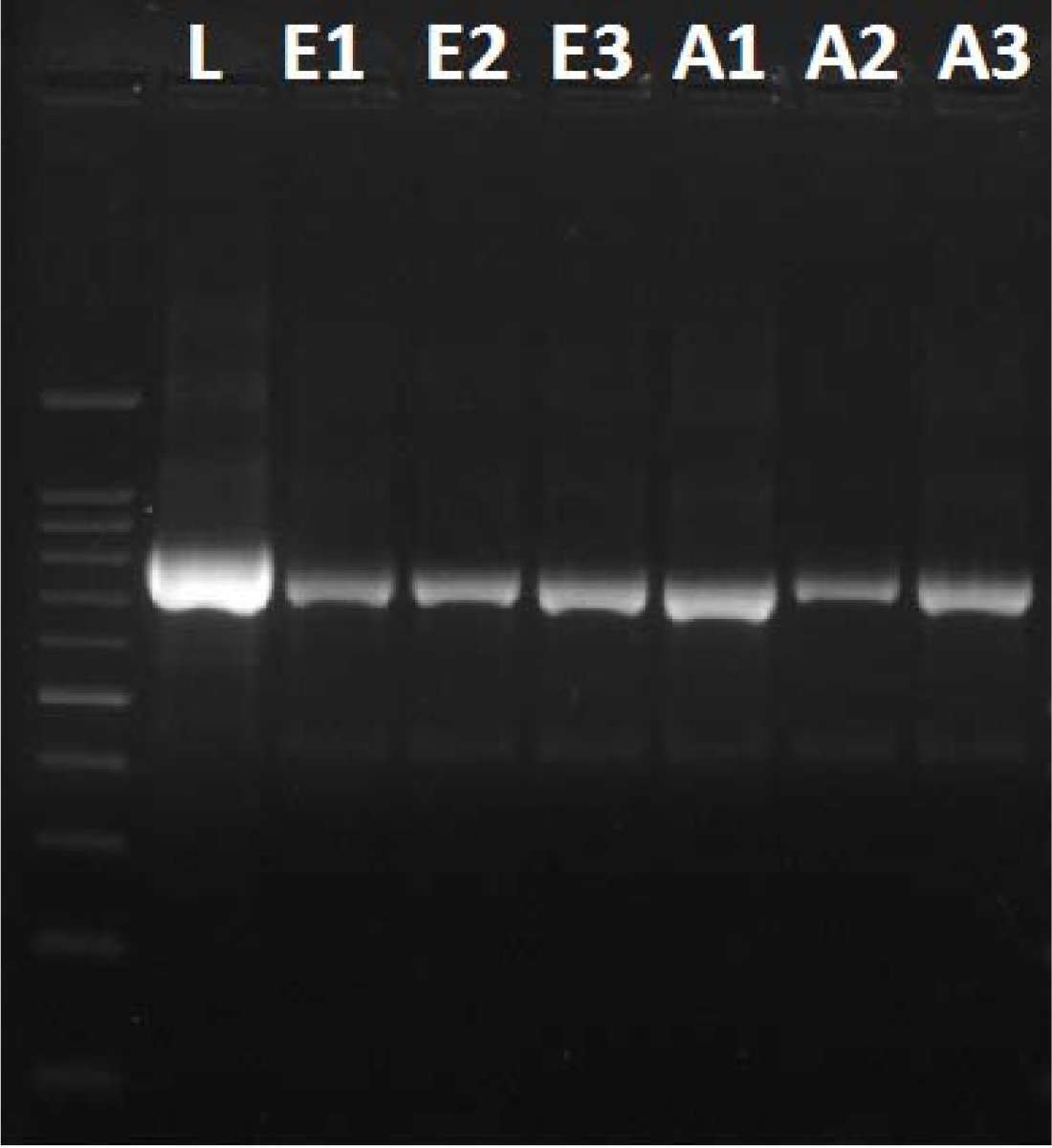
The PCR products of Rbcl DNA-Barcode for the Lab’s cotton (L) Egyptian (E1, E2, E3) and American (A1, A2, A3) varieties using the DNA isolated from fiber.

The NCBI blast of Rbcl band sequences results indicate that all DNA-barcoding technique was successfully to prove that the amplified DNA for all samples was actually cotton DNA though blasting to NCBI Database. The Rbcl PCR products digestion shows a different polymorphism between Egyptian and American cultivars (Figure 3). Dendrogram using Rbcl sequences shows that, Rbcl was almost successful to differentiate between local Egyptian cultivars and American imported cultivars (Figure 4).

**Figure 3:**
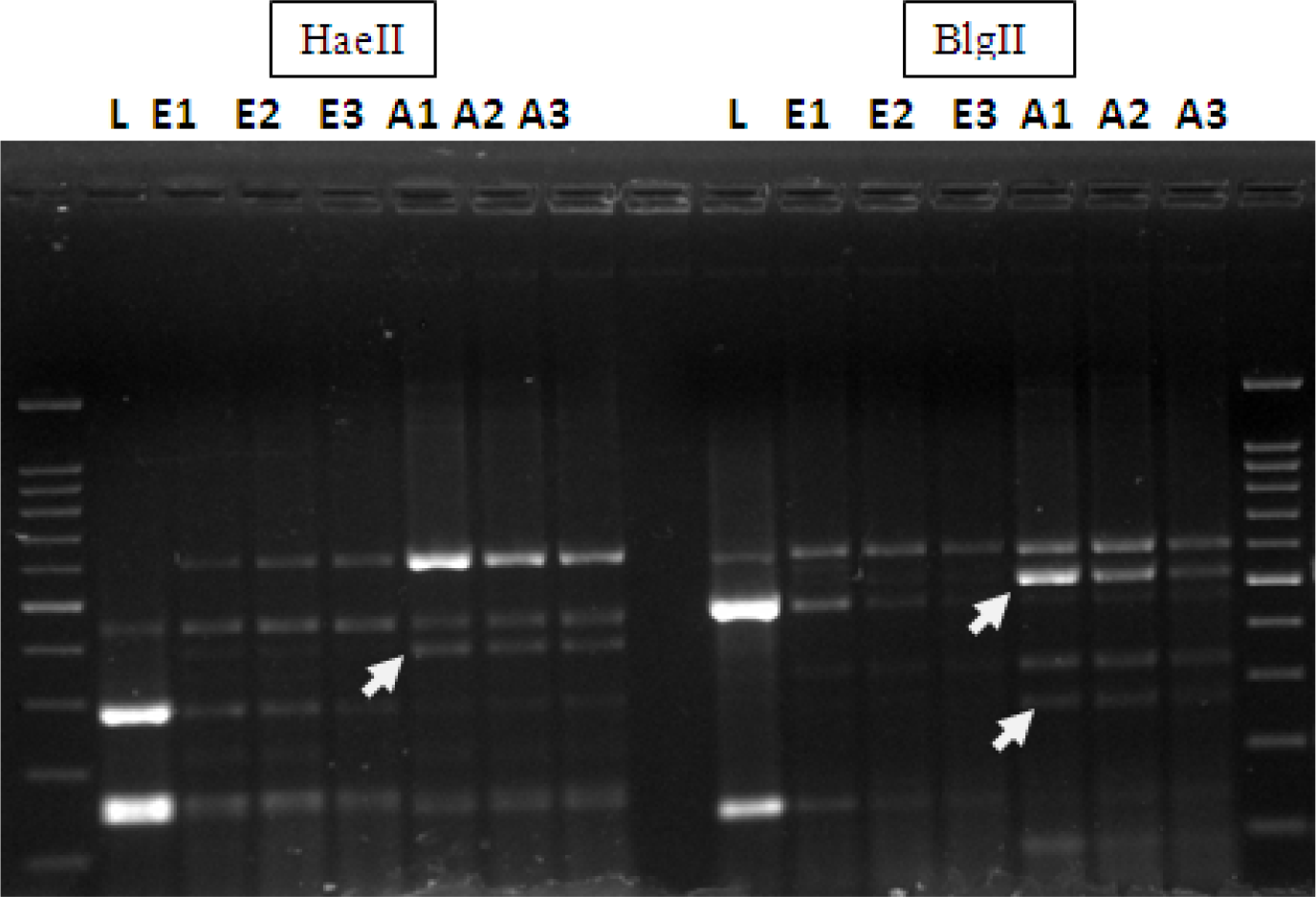
PCR amplification of Rbcl after cutting using HaeIII and BlgII enzymes, where **E1** to **E3** different Egyptian cotton varieties **A1** to **A3** are different American cotton varieties and the **L** is lab’s cotton. The white arrows show some bands differentiate between Egyptian and American genotypes:

**Figure 4:**
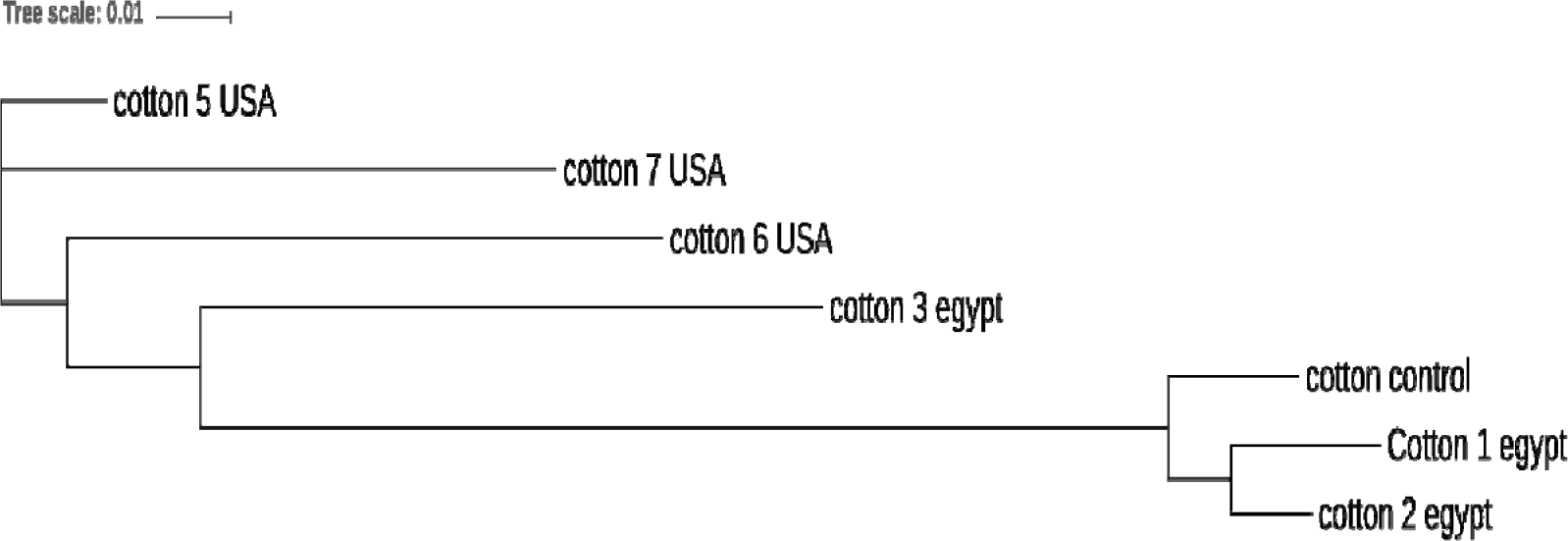
The sequence alignment using Rbcl sequences:

## Conclusions

Although the DNA isolation technique adds only the grinding with glass cover to CTAB protocol, it solved the problem of how we can get a proper amount of genomic DNA efficient enough for any PCR procedure from cotton fiber. The ability of this protocol to isolate DNA was validated even bleached and autoclaved cotton samples such as lab cotton fibers. The DNA-Barcoding assay helped indirectly to retrieve good PCR markers through CAPs with ability to differentiate between Egyptian and American. This is the first research article to report such results and we hope it could help Egypt and other countries import or export Egyptian cotton to confirm cotton textile origin.

## Acknowledgments

We thank the soul of Dr. Sami Adawy (MGGM lab) for his previous support to our lab during the past few years. We also thank Mr. Azmy Marzouk (MGGM lab) for helping.

